# Exploring social modulation: Microglia as a key mediator of individual immune response, plasticity and pathology in App-NL-G-F mouse model of Alzheimer’s disease

**DOI:** 10.1101/2024.01.23.576790

**Authors:** Fanny Ehret, Birte Doludda, Hang Liu, Sindi Nexhipi, Hao Huang, Warsha Barde, Fabian Rost, Rupert Overall, Andreas Dahl, Mirko HH Schmidt, Michael Sieweke, Gerd Kempermann

**Affiliations:** Institute of Anatomy, TU Dresden; DZNE – German Center for Neurodegenerative Diseases, Dresden; CRTD – Center for Regenerative Therapies, TU Dresden; DRESDEN-concept Genome Center, Center for Molecular and Cellular Bioengineering (CMCB) TU Dresden; Institute of Biology, Humboldt University, Berlin, Germany; OncoRay – National Center for Radiation Research in Oncology, Faculty of Medicine and University Hospital Carl Gustav Carus, Technische Universität Dresden; German Cancer Consortium (DKTK), Partner Site Dresden, and German Cancer Research Center (DKFZ); University of Bonn; Helmholtz-Zentrum Dresden - Rossendorf, Institute of Radiooncology – OncoRay

## Abstract

This study explores the influence of lifestyle on Alzheimer’s disease (AD) progression using App-NL-G-F mice in a complex enrichment system. Mice exhibited social deficits before plaque pathology or memory impairment, revealing a crucial link between lifestyle, behavior, and neuroinflammation. Plasma analysis indicates early inflammation and apoptosis-related changes, setting the stage for identifying markers predicting plaque manifestation. Beyond pathology, social behavior is linked to adult neurogenesis and microglia coverage, forming a dynamic connection with microglia activation. Further, sc-RNA sequencing unveiled a decrease in interferon-responsive microglia and alteration in antigen processing with enrichment. These findings underscore the beneficial impact of social housing on microglia and interconnected factors, pointing to microglia as a critical mediator of the behavior-pathology-plasticity interplay in AD. The study enhances our understanding of AD complexity and offers insights into potential therapeutic strategies, emphasizing the multifaceted nature of AD progression and the role of lifestyle in shaping its course.

## INTRODUCTION

Alzheimer’s disease (AD), a complex and multifaceted disorder, is shaped by the intricate interplay between genetic susceptibility and environmental factors. Addressing modifiable risk factors stands out as a promising avenue, potentially allowing for prevention or delay of up to 40% of dementia cases ^1^. While physical inactivity and low educational attainment are the highest morbidity risk factors^1, 2^, loneliness and social isolation have also been found relevant to AD progression particularly in preclinical stage^3^. The manifestation of AD exhibits remarkable variability, since a notable portion of this diversity can be attributed to lifestyle and behavior. Thereby recognizing the individual heterogeneity in intervention responses needs to be considered when investigating the underlying mechanisms in order to develop effective primary and secondary prevention strategies. Given that interventions during symptomatic stages might yield limited success, the importance of early lifestyle intervention becomes paramount.

We previously tackled certain aspects of behavioral intervention and identified abnormalities in a knockin animal model of AD^4^. Specifically, we observed deficits in habituation to an environment during presymptomatic stages, coupled with reduced stability in behavioral traits. Interestingly, residing in a complex enrichment environment not only enhanced inter-individual differences but also imparted a positive influence on metabolism. However, it is worth noting that within this initial study we refrained from further analyses of social interaction. This aspect became more relevant with the discovery that mice appeared prone to become disoriented and loose behavioral flexibility^4^, which possibly influence their social interaction. Social interaction contributes to a more robust cognitive reserve^5, 6^, which may act as a protective factor against the onset and progression of neurodegenerative diseases. Hence, active social life may thus be considered a valuable component for Alzheimer’s prevention.

Grasping the intricate interplay between behavioral activity and brain plasticity is a pivotal stride towards comprehending the intricate dynamics of neurobiology. Behavioral activity has been linked to brain plasticity, while adult hippocampal neurogenesis seems to be necessary to form stable behavioral trajectories ^7^and enhancing memory consolidation ^8-10^. Moreover, the plasticity influenced by adult neurogenesis, which is dependent on activity and experience, is linked to the individualization of the hippocampal circuitry ^11^. Consequently, enriched environment (ENR) not only amplifies phenotypic variability but also offers a valuable tool to explore gene-environment interactions ^12^. This sheds light on the importance of considering personal lifestyle factors in treatment strategies, an increasingly recognized aspect in the field of personalized medicine. To evaluate in our mouse model the impact of differences in individualization due to a genetic component, we will evaluate variance effects on adult neurogenesis as a measure of plasticity.

The interest in neuroinflammation as a modifiable feature of AD pathology has grown in conjunction with the genetic studies identifying AD risk variants. To what extent the immune system, as a key player for neuroinflammation in the course of AD, might be influenced by behavioral intervention was not clearly addressed in any long-term ENR experiment. While the role of innate and adaptive immunity in AD is relatively well-established, the exploration of behavior-related variation in immune factors and microglia responses as contributors to inter-individual differences in the progression of AD remains an uncharted territory in experimental research. Microglia, as the brains resident immune cells, are involved in immune surveillance and maintenance of homeostasis in the brain by phagocytotic clearance of pathogens, dead cells and protein aggregates. Further, recently conducted genome-wide association studies (GWAS) indicated that a large proportion of identified AD risk genes are enriched in microglia ^13, 14, 15^. Therefore, it remains to be clarified to which extent microglia show an individual response to changes in environment, activity or social interaction in healthy mice or conditions of AD predisposition.

To address these questions, we used *App* knock-in mice having Swedish, Bayreuth/Iberian and Artic mutations, with a mild onset in pathology starting at ∼3 months to evaluate the impact of environmental on presymptomatic stages. Mice were kept from 5 weeks to 7 months in a large ENR enclosure, consisting of 70 connected standard cages with various toys (houses, balls, nests, swings, tunnels, and bricks). The enclosure allowed for radiofrequency identification (RFID) tracking of individual mice within the cage system, down to the level of specific cages or tunnels they entered. Through longitudinal tracking, valuable insights into individual activity patterns and social interactions were gained. Initial immune cell response was evaluated from blood samples at age of 3 months. Microglia response and adult neurogenesis were evaluated by histological analysis at 7 months after 5 1/2 months of living in complex enrichment. In addition, single cell RNA sequencing of microglia was carried out from the different conditions, offering valuable insights into how ENR regulates microglia function in AD. Highlighting that social interaction and behavioral activity contribute to individual differences in immune and microglia response in AD – related neuropathogenesis.

## RESULTS

### Reduced social engagement and habituation behavior in NL-G-F mice at pre-symptomatic stages

Longitudinal analysis of NL-G-F mice, carrying the disease promoting Beyreuther/Iberian mutation and Artic mutation in addition to the Swedish Mutation, which alone does not have a functional phenotype, revealed differences in behavioral activity and social interaction. While subjected to complex enrichment from 5 weeks till 7 months of age (Fig. 1A), behavioral changes were analyzed at an individual level using RFID recording. This analysis aimed to calculate the individual exploration rate using mean RE, to compare behavioral trajectories of NL-G-F and NL mice. While the majority of NL mice habituate to the environment over time, as demonstrated previously ^4^, NL-G-F mice exhibit deficits in habituation behavior as time progresses (Fig 1B, C). In addition to these reduced adaptability traits in NL-G-F mice, social analysis reveals differences in their following behavior (Fig. 1F). Despite NL-G-F mice not exhibiting a genetic preference for whom they follow, as depicted in the pie chart, they do not decrease their following behavior over time to the same extent as controls. Instead, they distance themselves from others, leading to an increase in social distance over time (Fig. 1E). This pattern is also supported by social network analysis which is based on social distance measures (Fig. 1D). NL-G-F are characterized by reduced social engagement as they appear on the edges of the social network but in their social times, they follow more their counterparts, whereas controls get more individual over time, have more diverse social interaction and follow less. In conclusion, NL-G-F mice are social isolated and have difficulties to adapt to a changing environment with progression of pathology at pre-symptomatic stage, which might be as the earliest behavioral sign of AD.

**Fig. 1.**
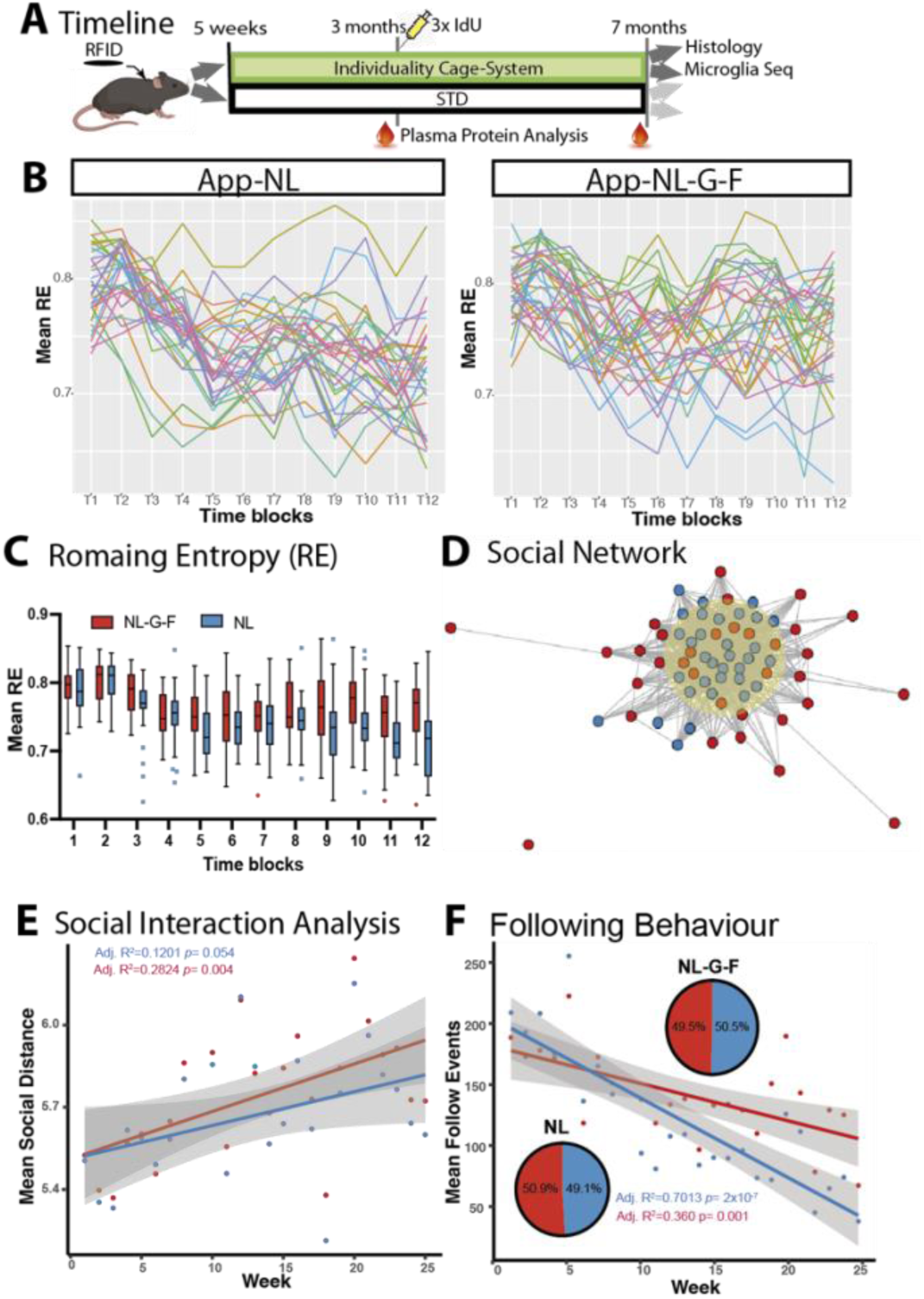
Deficits in social engagement and behavioral activity patterns in NL-G-F mice. **(A)** Experimental timeline outlines injections, entry into cage system, plasma collection and analysis parameter. **(B)** Individual behavioral trajectories recorded via RFID and plotted as mean RE over time for the different mouse lines. (**C)** Mean RE over time as box plots indicating that RE reduces over time in NL but not NL-G-F mice. Center line - median; upper and lower hinges - first and third quartiles; whiskers: Tukey. Repeated measures ANOVA with effect on genotype p> 0.0001, time and interaction p >0.0001 **(D)** Social network analysis on the basis of social distance depicts that more NL and less NL-G-F mice are in the center of the network. NL-G-F mice exhibited fewer social close contacts. **(E)** Social interaction analysis based on mean social distance over time, highlights the increasing social isolation of NL-G-F mice with time. Linear regression parameters are shown **(F)** Analysis of following behavior based on follow events of over time for each mouse line. NL-F-G did not adapt their behavior, whereas NL follow less at the end of the experiment. Linear regression parameters are shown in the graph. Pie charts depict genetic preference of following behavior; however, no preference was identified. ldU, Iododeoxyuridine; RFID, radiofrequency identification; RE, roaming entropy; STD, Standard housing; ENR, enriched environment,

### Unveiling the interplay of plasma proteins, behavioral activity, and plaque pathology

While looking for potential biomarkers of AD pathology, which can be influenced through environmental factor already at pre-symptomatic stages, we analyzed 96 proteins in plasma at 3 months (Fig. 2A). Our analysis revealed that 28 proteins are altered due to the environment, while 12 showed a genotype effect (Fig. 2B), and 7 proteins (Il1a, Casp3, Il23r, ccl20, Erbb4, Cntn1, Cdh6) displayed a genotype-environment interaction (Fig. 2C, D). The proteins that were identified are expressed and relevant for a multitude of organs beyond the central nervous system (Fig. 2C), playing a central role in receptor signaling pathway, inflammation, apoptosis and neuronal differentiation, as evaluated by string analysis (Suppl. 1). We conducted further analysis to explore potential correlations with pathology or lifestyle among the analyzed markers (Suppl 1). We specifically investigated two candidate proteins, Casp3 and Il1a, which exhibited a favorable impact of enrichment at 3 months (Fig. 2E, F). Subsequently, we also assessed these proteins in plasma samples at the end of the experiment (Fig 2G, H); however, their positive regulation was no longer evident at 7 months. Nevertheless, Casp3 in plasma, as a mediator of apoptosis, correlated with behavioral exploration at 3 months (Fig. 2I), but not at 7 months. Similarly, Il1a, an inflammation-related cytokine, displayed a contrasting activity-depend correlation pattern between controls and AD-like mice (Fig. 2J).

**Fig. 2.**
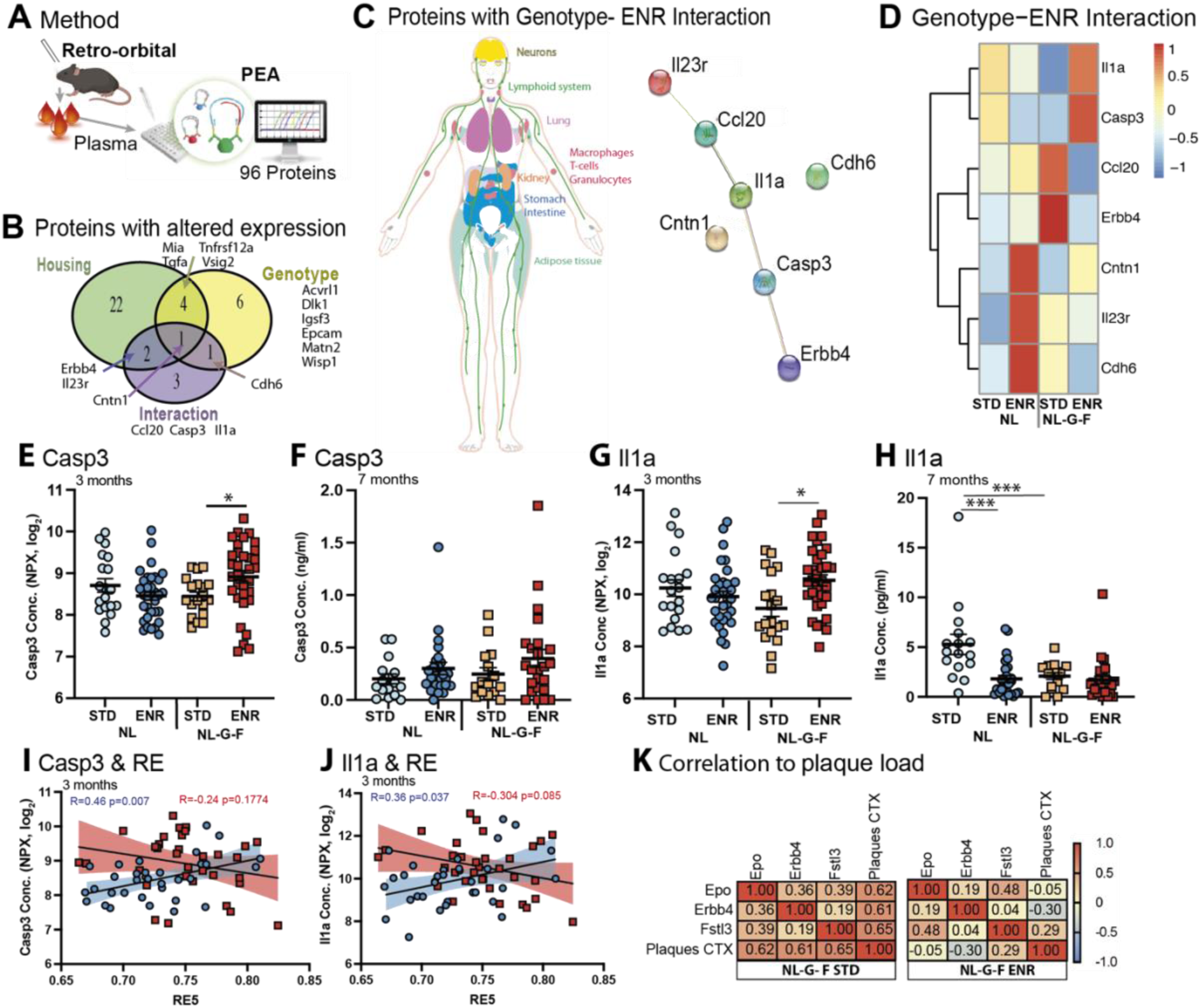
Analysis of plasma at 3 months identifies proteins relevant for inflammation, apoptosis and plaque pathology. **(A)** Illustration of the method used to isolate and analyze plasma samples at 3 months, after 8 weeks of enrichment. **(B)** Venn diagram highlighting proteins with a significant effect for the different parameters (housing, genotype and interaction of both).(**C**) Illustrates the expression of the 8 proteins with significant genotype ENR interaction across the human body (**D**) Spearman correlation matrix with hierarchical clustering of the proteins with Interaction effect, depicting relative expression in the different conditions. (**E**) Protein expression of Caspase 3(Casp3) at 3 months measured by PEA revealed a significant interaction effect, 2way ANOVA F_(1,100)_= 6.718 p=0.011, with STD vs ENR in NL-G-F, *p=0.036*; N= 19 for each STD cohort and N=33 for each ENR cohort on all PEA assays (**F**) Casp3 levels measured by ELISA (**G**) Concentration of Il1a measured by PEA at 3 months with a significant interaction effect F_(1,100)_=6.706, *p= 0.0079*, with STD vs ENR in NL-G-F, *p=0.025*. (**H**) Il1a concentration at 7 months measured by ELISA did reveal a significant effects of genotype F_(1,84)_=9.829 p=0.0024, housing F_(1,84)_= 13,54 *p=0.0004* and interaction F(1,84)= 8.505 p=0.0045; with STD vs ENR in NL p<0.0001 and STD NL vs STD NL-G-F p<0.0001.(**I,J**) Correlations analysis of Casp3 concentration (**I**) and Il1a concentration (**J**) at 3 months in relation to roaming entropy (RE) of the corresponding time block ( 10 days). Pearson’s R for the different lines are depicted. (**K**) Correlation matrix of all regulated proteins with significant effects to plaque load in the cortex region as depicted by color code. Values plotted represent Pearsons R. Significant interactions are shown as *p< .05, **p < .01, ****p < .0001. ELISA, enzyme-linked immunosorbent assay; PEA, proximity extension assay; Con., concentration; CTX, cortex; STD, standard housing; ENR, enriched environment.

Although none of the investigated proteins showed a regulatory response to social parameters, we did identify a protein (Erbb4) that exhibited a correlative response to pathology (Fig. 2K) and behavior (RE5) within the subset of proteins with genotype housing interaction (Suppl. 1 C-D). Furthermore, two proteins (Epo, Fstl3) also correlated with plaque pathology; however, only a regulatory response to enrichment, not genotype, was observed. Notably, these three proteins displayed a significant positive correlation with cortex plaque load in NL-G-F mice housed under standard conditions. However, this correlation weakened and even displayed opposing trends under the enrichment condition (Fig. 2K), underscoring their potential early relevance in the context of AD pathology.

### Individual activity patterns and social parameters in adult neurogenesis and their association with AD pathology

To identify alterations and establish a link between activity patterns in adult neurogenesis and AD pathology, we administered IdU to mice at 13 weeks, following 8 weeks of enriched housing. The number of IdU+ cells was enhanced in NL and NL-G-F mice under ENR conditions compared to STD housing (Fig. 3A). However, this enhancement was slightly lower in NL-G-F mice. At this stage, we were unable to find a correlation with individual following behavior or social factors (Fig. 3B), possibly due to relative short run-time.

**Fig. 3.**
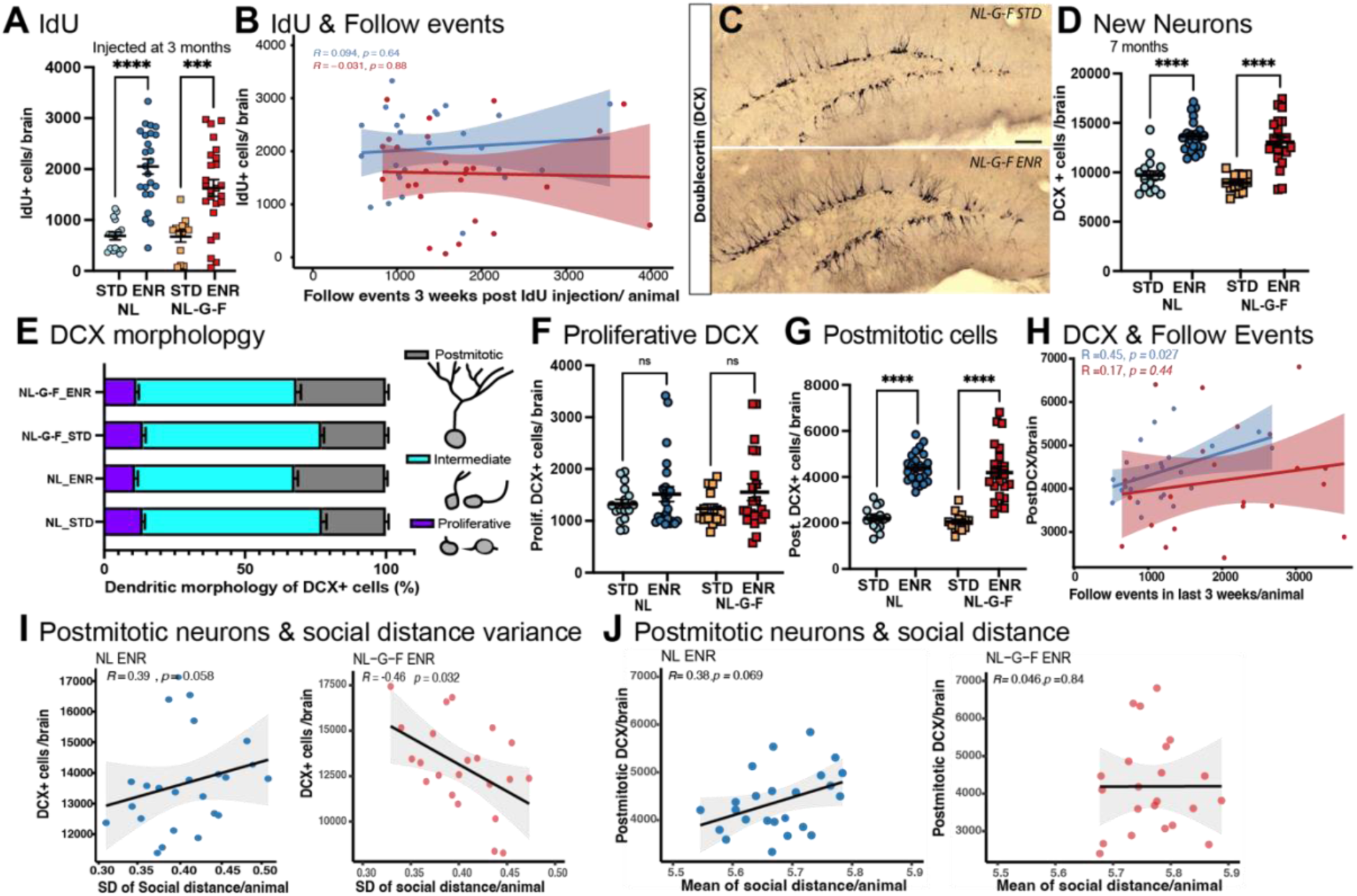
Adult neurogenesis in the hippocampus correlates to aspects of social behavior. (**A**) IdU, an injectable tracing marker, used to analyze neurogenesis at 3month. IdU+ cells in hippocampus were increased after ENR housing, two-way ANOVA F_(1,77)_ = 57.81 *p<0.0001*. Further, variance was increased in both genotypes after ENR, as measured by Brown-Forthsythe *p<0.0001*. (**B**) At this stage no correlation to behavior, as presented by Pearson correlation. (**C**) Micrographs depicting the difference in Doublecortin (DCX) expression in STD vs. ENR housed NL-G-F mice. Scale bar 100µm. (**D**) The number of DCX positive cell analyzed at 7 months are higher in ENR in comparison to STD housing, two-way ANOVA F_(1,73)_ = 90.85. Effect on variance STD vs ENR Brown-Forthsythe *p<0.0001* (**E**)Classification of DCX+ cells based on morphology into proliferative, intermediate and postmitotic state. For each condition the cell states were plotted as stacked bar graph. (**F, G**) The total cell number of DCX+ cells were evaluated in the morphologies, with no effect on cell proliferation stages but on postmitotic stages due to ENR F_(1,73)_= 127.3 *p<0.0001*. (H) Postmitotic DCX are correlated to the following behavior in the last 3 weeks of ENR, controls but not NL-G-F, showed significant interaction as shown by Pearsons R. (**I**) The interaction of individual standard deviation (SD) for social distance over the entire ENR period and DCX+ cells in hippocampus for each animal are presented. Spearman’s R are presented in the graph. (**J**) The mean of social distance in correlation to the number of postmitotic DCX cells., Spearman’s R are shown. Significant interactions are shown as \*\*\**p < .001;* ns, non-significant. STD, Standard housing; ENR, enriched environment, IdU, iododeoxyuridine.

Further analysis focused on evaluating newly generated neurons using DCX at the end of the 7-month experiment (Fig. 3C). Although, we did not observe any genotype-related deficits in DCX+ cell counts; nevertheless, a notably positive and robust impact due to ENR persisted even at this early pathological stage. We further classified these cells based on their morphology into proliferative, intermediate, and postmitotic categories, aiming to identify differentiation-related changes (Fig. 3E-G). ENR appeared to positively influence postmitotic neurons even under pathological conditions. Additionally, we identified significant interplays with social parameters. The number of postmitotic DCX cells exhibited correlation with following behavior in the final 3 weeks of enrichment. In control mice, an increased tendency to follow others correlated with higher rates of adult neurogenesis. However, this relationship was disturbed in NL-G-F mice (Fig. 3H). Further, control animals showed an increase in variance in relation to their distance with other mice in the system, resulting in higher intra-individual variance (higher SD, Fig 3I) and had more DCX+ cells, whereas the opposite was true for NL-G-F mice. This suggested that the stable interactions (less variability) and reduced adaptability in NL-G-F mice resulted in a more newly generated neuron (Fig. 3I). In addition, the higher their social distance in controls– reflecting more following behavior and more dominant related-behaviors– the greater the number of new neurons observed, a trend not evident in NL-G-F mice (Fig. 3J). Hence, situations involving more novel and varied exposure require flexibility, an aspect that is already altered in NL-G-F mice, Thus, the stability and following behavior play a more dominant role, counteracting natural exploration. In conclusion, our findings underscore the change of social interaction and activity on adult neurogenesis in early stages of AD. Subsequent analysis will delve into whether microglia cells mediate this regulatory phenomenon.

### Behavioral exploration and social interaction: shaping microglia morphology and activation, and their correlations with adult neurogenesis

We performed semi-automated analysis to examine microglia (Iba1) morphology, proliferation (Ki67), and activation (Trem2) in 3-4 histological sections per animal from hippocampus and cortex (Fig. 4E, L). The Fiji code used for automatic segmentation was first validated in test samples (Suppl. 2). Since exploration and social engagement is different in our mouse model it was remarkable to find that independent of genotype there is a direct relationship between social distance and the microglia coverage in the hippocampus (Fig. 4A) but not the cortex (Suppl. 2). Thus, the greater the social interaction among mice, the higher the number of microglia detected, highlighting the significance of hippocampal microglia involving in social interaction possibly by mediating the process of adult neurogenesis. In neurogenerative conditions microglia transition from a homeostatic state into a disease associated microglia (DAM) state, with an increase in soma size, amoeboid morphology and reduced ramification. Further analysis of morphology revealed, NL-G-F mice displayed larger soma sizes compared to NL mice regardless of housing conditions (Fig. 4B), which is also true for the avg. M-score (Fig. 4C). The M-score assesses microglia ramification based on the area and circularity of individual Iba1+ cells. A smaller M-score is attributed to less round and smaller cells (indicative of resting, ramified microglia), while a larger M-score results from more round and larger cells (reflecting phagocytic, amoeboid microglia). Notably, a significant correlation emerged between M-score and cumulative RE in NL mice, while this was not observed in NL-G-F mice (Fig. 4D). This indicates that the impact of behavioral activity on morphology is disbalanced under AD conditions.

**Fig. 4.**
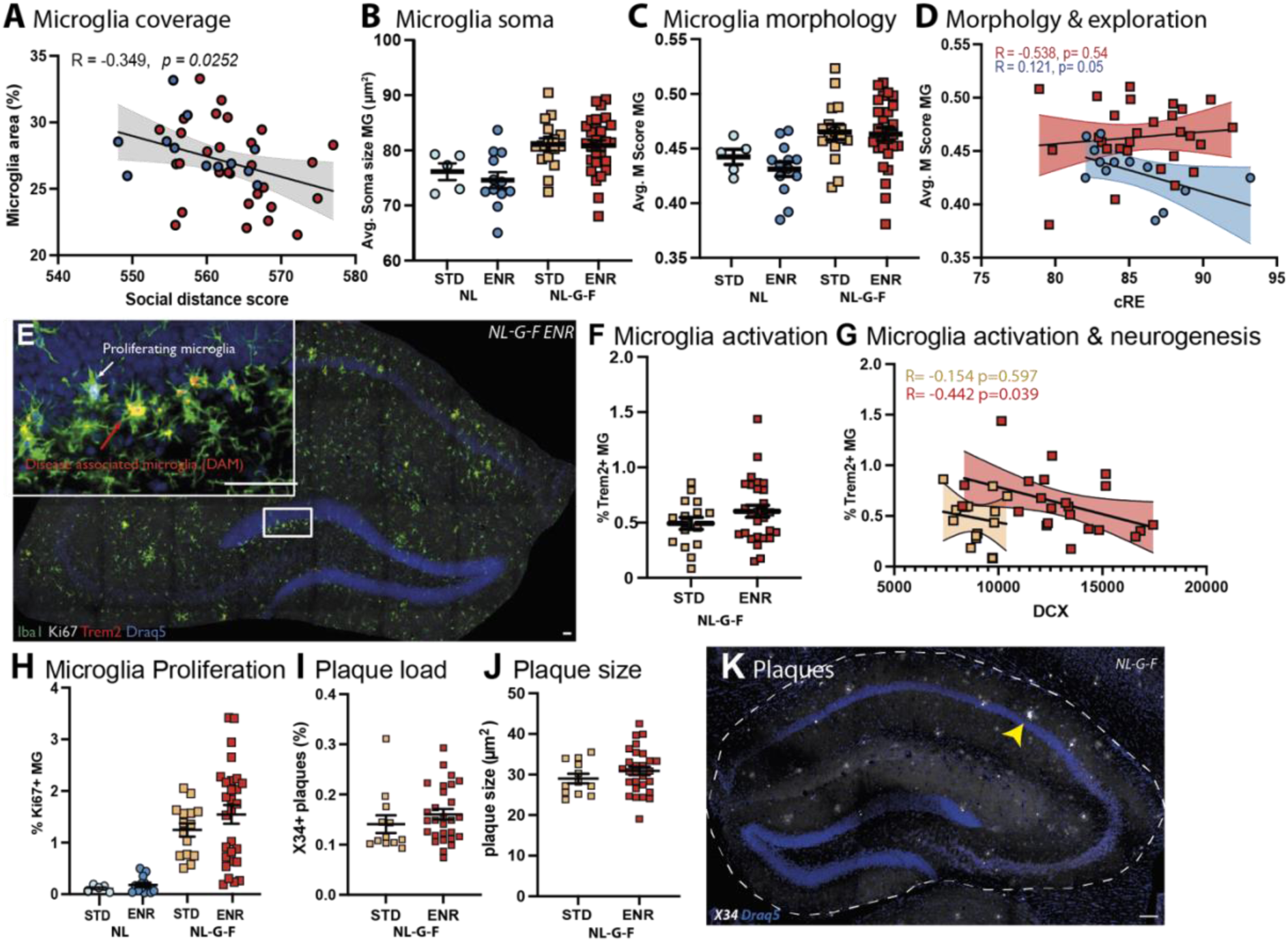
Histological microglia and plaque analysis in hippocampus implicate microglia as mediator of environment x plasticity interaction. **(A)** Automated segmentation of microglia from Iba1+ cells revealed correlation to social distance independent of genotype, Pearson’s R is presented in graph **(B**) An effect on soma size due to genotype was detected by two-way ANOVA F _(1,58)_ = 14.22 NL vs NL-G-F *p= 0.0004*. (**C**) Analysis of changes in microglia morphology by M-score (as a measure of ramification) is in compliance to soma size and showed an effect of genotype but not housing condition two-ANOVA F_(1,58)_ = 10.02, NL vs NG-G-F *p= 0.0025*. (**D**) Correlation of cumulative explorative behavior (cRE) as a longitudinal measure of activity and M-score, A significant relation in controls is shown by Pearsons R. **(E)** Representative micrograph used for automated segmentation of microglia based on Iba1, proliferation marker Ki67, activation marker Trem2. (**F**) disease activation of microglia by Trem2 expression was not influenced by the different housing conditions. (**H**) The percentage of Ki67+ microglial cells, as a measure of proliferation, is increased in NL-G-F in comparison to NL, two-way ANOVA F_(1,58)_ = 32.79, *p<0.0001* and further an increase variance was seen upon ENR, as evaluated by Brown-Forsythe test *p<0.0001*. (**I-K**) Plaque load and size in the hippocampus was analyzed by X34, a derivate of Congo red. (**K**) Representative micrograph used for segmentation of plaques, a large dense plaque is marked with yellow arrow. Plaque coverage (**I**) nor plaque size (**J**) was not influences by housing condition. ENR, enriched housing; MG, microglia; STD, standard housing, cRE, cumulative roaming entropy. Scale bar in E, K 50µm.

In addition, microglia proliferation was observed to be higher in NL-G-F mice than NL mice, but ENR did not significantly increase proliferation; instead, it amplified variance (Fig. 4H). Although no significant difference in Trem2-dependent activation was found between STD and ENR conditions (Fig. 4F), a noteworthy connection emerged between adult neurogenesis and Trem2 activation, particularly in ENR (Fig. 4G). This connection implies that higher inflammation and Trem2 activation correspond to lower numbers of newly formed neurons. Since microglia are modulators of neurogenesis by phagocytosing apoptotic cells ^16^a shift to more Trem-2 activation and thus involvement in inflammatory response might negatively impact modulatory function for adult neurogenesis in AD conditions. Since analysis was performed during early stages of pathology no significant impairment could be detected. In addition, in the hippocampus region, plaque levels and microglia activation seem to be interconnected. Plaque levels were assessed via immunohistology using X34, a fluorescent derivative of Congo red (Fig. 4K). Despite similar plaque levels and sizes between STD and ENR condition (Fig. 4 I, J), a correlation with microglia activation was identified under STD conditions but not in ENR in the hippocampus (Suppl 2D). The cortex region was evaluated, but no significant correlations of this nature were observed. NL-G-F mice in ENR showed trends towards reduced average soma size (Suppl 2I), while no notable impact was observed on proliferation or Trem2-dependent activation (Suppl 2 K-L). This suggests that Trem2-dependent activation might represent fundamental programs less influenced by behavior but with critical impact on neuroinflammation. Nonetheless, ENR appears to positively influence individual differences, as evidenced by the variance effect on proliferation in both regions. Taken together this highlights that microglia might play a mediatory role for individual immune response by adjusting morphology, proliferation and activation in response to social and behavioral activity and pathology.

### Microglia response to enriched housing: Identification of cellular and molecular changes

To provide a comprehensive understanding of the influential effect of complex enrichment on cellular and molecular signatures of hippocampal microglia cells at 7 months from this cohort, we conducted single cell RNA sequencing (sc-RNA-seq) of FACS sorted Microglia cells (Ly6C^-^ CD45^low^ CD11b^+^), using Clec7a/Dectin-1 as an activation marker (Fig 5A) detecting cells in a proinflammatory state in neurodegeneration ^17^. Consistent with the histological findings, microglial activation was evident exclusively in NL-G-F mice, displaying little variations across the two housing conditions. Moreover, a diminished microglial presence in NL mice compared to their NL-G-F counterparts was observed, aligning with the histological dataset. From the 16 mice used for hippocampal microglia isolation, 5823 cells could be used for scRNA transcriptome analysis after quality control and normalization. The application of unsupervised Leiden clustering to the scRNA seq data led to the identification of seven distinct subclusters, each characterized by unique gene expression profiles (Fig.5B, 6A). The assignment of these clusters to cellular phenotypes was based on established gene expression profiles, primarily drawn from previous publications of different mouse models ^18, 19^ (Fig. 5C). Compositional analysis (Fig. 5B) unveiled a reduction in homeostatic phenotypes (0-1) in NL-G-F mice, accompanied by a shift towards DAM activation stages (2, 4).

**Fig. 5.**
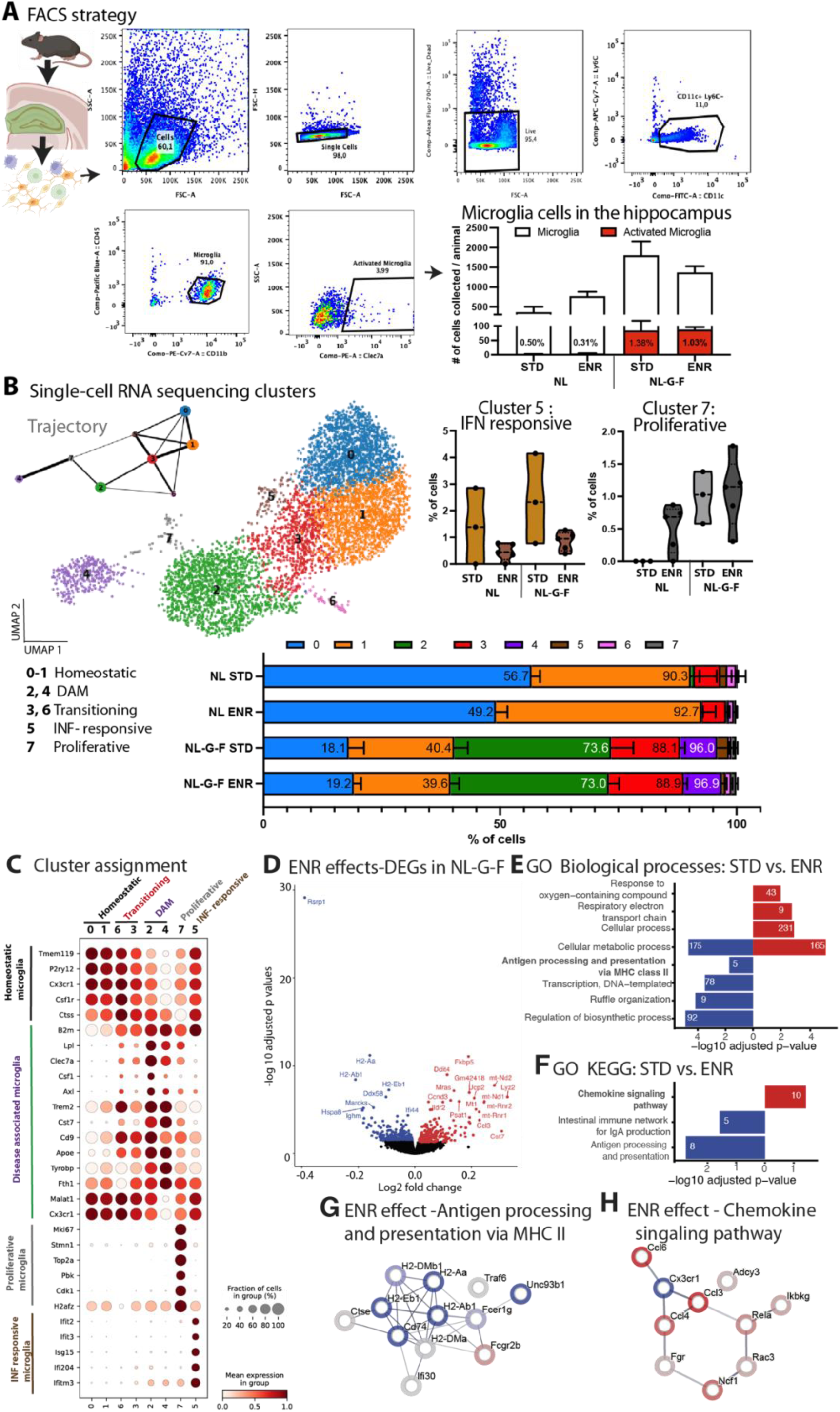
Single cell RNA sequencing of hippocampal microglia highlights cellular shifts and gene expression changes upon ENR. (**A)** Illustration of the applied method and used gating strategy to isolate microglia from aged hippocampal tissue. As selection marker were used CD11c, Ly6C, CD45, CD11b and Clec7a just for quantification of activation state. The percentage of activated microglia cells in the different conditions is stated in the bar graph. (**B**) Sequencing of hippocampal microglia from the different conditions (from each genotype N=5 for ENR, N=3 from STD condition). In total 5823 cells were analyzed by unsupervised Leiden clustering. Seven different clusters were identified as shown on UMAP. The assignment of the clusters to cell states, as shown on lower left, was verified by marker genes, see **C**. Trajectory analysis (top left) of the identified clusters using PAGA to observe the continuum of state changes within this dataset. The proportions of cells in the identified clusters from the different conditions are shown as stacked-bar graph. The small clusters 5 (interferon-responsive microglia) and 7 (proliferative microglia) showed a relevant regulation upon ENR. (**C**) Dot plot depicting gene expression in the different clusters, the cluster number assigned are shown at the top. Each row represents the level of expression of a selected key gene. The color represents the scaled expression changes. Size of the dot represents the fraction of cells in a given cluster (**D**) Volcano plot of the ENR effect in NL-G-F mice on DEGs, significant hits were color coded for down regulation (blue) and up regulation (red). (**E-F**) Identified DEGs were analyzed by Gene ontology (GO) for biological processes (**E**) and KEGG pathways (**F**). Selected significant terms were plotted as bar graph, color coded for down (blue) and up (red) regulated processes or pathways, the number of genes significantly altered are presented inside the bar. (**G -H**) Using STRING with ranks, two relevant protein networks identified in **E** and **F** were mapped with color coded halo based on log-fold change (FC).) Edges represent protein-protein interaction. DAM, disease associated microglia; FACS, fluorescence activated cell sorting; INF, interferon; ENR, enrichment; KEGG, Kyoto encyclopedia of genes and genomes; MHC, major histocompatibility complex; INF, interferon.

Two clusters are of critical importance: First the proliferative cluster (7) validated the histological data and affirmed the heightened proliferative capacity in App-NL-G-F and increased variance in following ENR in both genotypes. Second, interferon (INF)-responsive microglia (5) displayed susceptibility to housing conditions in ENR, pointing towards a reduction of their inflammatory microenvironment, even in the NL background. These observations indicate that while DAM activation appears unaffected by ENR, an positive impact on distinct clusters and inflammatory activation can be seen.

Trajectory inference predicts that the majority of proliferative cells derive from or transition into DAM clusters, which develop primarily from homeostatic and transitioning microglia as reported previously ^18^. INF-responsive microglia, however, do not transition directly from DAM clusters, but rather from homeostatic microglia.

Analysis of gene expression changes in the different conditions revealed intriguing targets in NL-G-F mice (Fig. 5D-H), while a focused examination of cluster 5 or 7 was infeasible due to low cell numbers, one target Ifi204 was significantly enhanced after ENR in cluster 5 (adjusted *p = 0.023*) which plays an important function in innate immunity^20 21^ since its relevant for inflammatory cytokine production (DEGs are provided as Suppl. table). Subsequently we shifted our focus to differentially expressed genes (DEGs) in the STD vs. ENR in all cells (Fig. 5D) and by using gene set enrichment analysis gene ontology (GO) topics were identified with significant alterations in biological processes (Fig. 5E). Notably, ENR exhibited a favorable influence on genes relevant for the response to oxygen compounds, respiratory electron transport chain, and metabolic processes. Several genes relevant to cellular metabolic processes displayed elevated or reduced expression, likely due to their distinct functions. Remarkably, ENR led to reduced gene expression relevant for antigen processing and presentation. Given their high relevance for microglia function and activation, a detailed analysis of the altered genes interaction network and its effect size are depicted (Fig. 5G). Further, DNA-transcription, ruffle organization and regulation of biosynthetic processes seemed diminished. A KEGG pathway enrichment analysis (Fig. 5F) highlighted the positive regulation of chemokine signaling in NL-G-F mice after ENR, alongside a reduction in intestinal immune network-related genes for IgA and further implications for alterations in antigen processing and presentation. To delve deeper into chemokine signaling of microglia, we mapped the involved genes and color-coded the effect size accordingly (Fig. 5H). Thus, although microglia’s acute activation stage does not undergo significant change post ENR, several signaling pathways show positive effects, indicating a central role for microglia in AD immune response that is favorably influenced by enrichment and social engagement.

## DISCUSSION

This study emphasizes the critical influence of emerging individual heterogeneity and the profound impact that social interaction and behavioral exploration have on central and peripheral immune response as well as brain plasticity possibly mediated by microglia in AD. We propose that microglia play a pivotal mediatory role in shaping behavioral responses within the context of the intricated interplay between plasticity and pathology. Notably, this study is the first to reveal that AD-like mice, even at presymptomatic stages, exhibit deficits in social parameters prior to the onset of pathology or memory impairment. NL-G-F mice, as previously discussed, not only display reduced adaptability but also had less close social connections over time. This deficit exerts a significant influence on hippocampal plasticity and microglial coverage. Additionally, the peripheral immune profiles show differential alterations in response to ENR in AD-like mice, featuring changes in proteins with known relevance to Alzheimer’s disease and cytokine receptor binding, even at presymptomatic stages. While a direct link between microglial response and peripheral immune response was not fully established, a correlation to plaque load could be identified, highlighting a need for further investigations of presymptomatic alterations. Besides the change in INF-responsive microglia population, our microglia sequencing data revealed an improvement of chemokine signaling as well as antigen processing and presentation pathways upon ENR in AD-like mice. In essence, these novel observations provide first insights into the complex interplay between microglia, plasticity, and immune cell activation, shedding light on the wide spectrum of individual variances in this context.

To the best of our knowledge, there has been limited exploration of preclinical behavioral abnormalities in AD. While some studies have hinted at a link between loneliness and increased amyloid burden in cognitively normal older adults, there is a significant gap in our understanding of social deficits and social networks in individuals during the preclinical stages of AD. Although research in various AD mouse models has shown that social deficits and anxiety can manifest in middle-aged and older mice ^22-24^, it’s crucial to emphasize that very few studies, including our prior work, have specifically addressed altered behaviors, such as changes in exploration, rearing, and social memory, in mice exhibiting AD-like characteristics during the pre-symptomatic stages ^4, 23, 25, 26^. In our current study we were able delved deeper into the aspect of social structure and following behavior over time due large cohort and longitudinal analysis over 5 1/2 months. Our analysis revealed a decline in following behavior over time in both genotypes, indicating the establishment of a stable hierarchy. A parallel study by Doludda et al. ^27^ highlighted that following behavior can reflect dominance-based behavior. Hence, when following behavior decreases over time, it suggests that less dominance-related behavior is needed to maintain the social hierarchy. In essence, the App-NL-G-F mice in our study had to reaffirm their hierarchy position and struggled to establish a close social network, reflecting a situation akin to that seen in patients. As disease progression advances, patients often experience significant social impairment, which remains a challenging burden and affect to a great extent all patients outcomes.

In the context of measuring brain plasticity through adult neurogenesis, intriguing connections to social parameters have been identified. NL-G-F and control mice exhibit contrasting interactions, revealing a noteworthy association. In controls, numerous social interactions with divers’ partners and selected following behaviors enhance brain plasticity, leading to higher neurogenesis rates. This suggests that those engaging in more dominant-related behavior and overall social isolation experience greater neurogenesis, supporting the notion of hierarchy positions influencing hippocampal neurogenesis ^28^. The opposite is true for NL-G-F, were more stable social interactions with less partners and higher ratio of following behavior resulted in higher neurogenesis rates. Thus, its most likely that AD-like mice face challenges in adapting to ever-changing social interactions and social dominance, preferring constant contact with a small group of mice. Despite these struggles, maintaining social interaction and the associated sensory, cognitive, and physical stimulation is crucial for preserving brain plasticity as long as possible in the context of AD manifestation, even in rodents.

Microglia likely play a crucial mediatory role in the intricate interplay between plasticity x pathology in AD. Recent research has offered new insights by linking variations in microglial morphological profiles to individual behavioral and emotional characteristics^29^. Consistently, we now observed a negative correlation between the M-score of hippocampal microglia and cumulative RE values, which implies that mice with higher exploratory ability have less activated microglia. Further, the more social animals are the more microglia coverage was seen in the hippocampus, independent of genotype. However, the number of proliferative and activated microglia was not impacted by behavioral or social parameters when evaluated at the end of ENR housing. As a support of the hypothesis that the communication between peripheral immune system and central nervous system assists behavioral responses to the environment ^30^, the level of Il1a as a peripheral pro-inflammatory cytokine was altered in NL-G-F and could be normalized ENR conditions at early stages but not at 7 months. In alignment, alteration in plasma levels at 3 months reflect the human early-moderate stages and cannot be compared to the inflammatory situation at late stages^31^, where increased inflammatory markers are related to a higher risk of dementia. Further, Erbb4 was found as a candidate, which showed a connection to cortical plaque load. Erbb4 is known in its function in migration, differentiation and apoptosis and has recently been shown to mediate Aß-induced neuropathology ^32^. Errb4 is increased in NL-G-F mice and can be significantly reduced by ENR to control levels, highlighting its positive impact.

To further discuss the critical role of microglia in AD conditions, one should point out that microglia also serve a phagocytic function in adult neurogenesis to clear the majority of newborn neural progenitors and help maintain the equilibrium of the neurogenic niche ^33^. Thus, overactive microglial in neurological disorders might inhibit adult hippocampal neurogenesis^34^. The upregulation of Trem2 with advancing amyloidosis promotes microglia to acquire a phagocytic phenotype, which explains the negative correlation between the number of Trem2+ microglia and newborn neurons. Although the proportion of cells in the two DAM stages are not differentially impacted by ENR, analysis of differentially expressed genes upon ENR housing revealed a down regulation of MHCII related genes, pointing towards a response onto the delicate interplay of neuronal, microglia and Immune cells in AD. As shown previously, microglia can act as antigen-presenting cells via MHC II to CD4+ T-cells ^35^ eliciting adaptive activation and T-cell clonal proliferation, which stimulates adaptive immune response in CNS. As recently pointed out, Antigen processing and chemotaxis were critically upregulated along the Aß and genotype axis in aged mice^36^. We have now demonstrated that ENR has immunomodulatory functions by reducing gene expression relevant to antigen processing and presentation via the MHCII pathway and enhancing proinflammatory chemokine signaling (Ccl3, Ccl4, Ccl6). This is crucial for the crosstalk of central to peripheral immune communication^37^ and may also impact astroglia crosstalk in CNS immunity. The observed downregulation of Ighm expression following ENR, a critical component in the NFkB glia inflammasome pathway, implies modifications in these pathways that could influence the inflammatory response of microglia. These changes potentially have implications for disease manifestation and T-cell infiltration in later stages and thus greatly impact the progression of the disease. Moreover, we observed a decrease in INF-responsive microglia following ENR housing. Interferon signaling is a pivotal regulator of microglia phenotypes and neuroinflammation. Specifically, type I INFs, named for their ability to "interfere" with viral replication, have been identified to play distinct roles beyond infection, with an increase observed in AD pathogenesis^38^ The current speculation revolves around the idea that the type I INF system represents signaling pathways with a modifiable role in AD progression^39, 40^. Type I INFs, capable of positively regulating of interleukins like IL1ß, IL6 and TNFα. While, blocking of type-I INF signaling, reduced microglia activation and phagocytosis^41^. Therefore, the nuanced regulation influenced by ENR has a positive impact on this microglia population, effectively reducing these cells back to control levels.

The observed alterations in microglia profiles, coupled with the impact on neuroinflammation, emphasize the potential of environmental enrichment to modulate key players in AD pathology. These findings not only contribute to our understanding of the complex dynamics underlying AD progression but also pave the way for targeted interventions that harness the beneficial effects of lifestyle modifications in mitigating disease-associated changes.

## METHODS

### Animal husbandry

App ^NL-G-F/NL-G-F^ and App ^NL/NL^ mice ware obtained from Rinken Institute containing a Swedish (KM670/671NL), Arctic (E693G) and Beyreuther/Iberian (I716F) mutation, as described previously ^42^. All 104 used female animals from the 2 mouse lines were maintained on a 12 h light/dark cycle with food and water provided *ad libitum* at the DZNE, Dresden, Germany. Control and enriched animals received the same fortified chow (no. V1534; Sniff) with 9% of energy from fat, 24% from protein, and 67% from carbohydrates. The experiment was licensed from the local authority (Regierungspräsidium Dresden) under 25-5131/496/31. At the age of 4 weeks, half of the animals were randomly assigned to the two different housing conditions either living in custom-built system of enrichment (PhenoSys GmbH, now marketed as “PhenoSys ColonyRack Individuality 3.0”) referred to as enriched housing (ENR) or standard housing (STD). The experimental groups consisted of 33 female animals in ENR per genotype, housing 66 animals together in the same enriched environment. Further, 19 female animals per genotype in STD condition, housed separated for each genotype. At 5 weeks of age, animals were subcutaneously injected into their neck with a glass-coated microtransponders (SID 102/A/2; Euro I.D.) under brief isoflurane anesthesia. At 6 weeks, mice entered into the different housing conditions. In the ColonyRack the environment changed over time, toys, nesting material and huts chanced in position and complexity biweekly. Food and water locations remained the same over the entire period. After 13 weeks, mice were injected with 3x IdU (57,5 mg/kg) and blood samples were collected from vena facialis in the cheek. After 28 weeks of ENR, all mice left the cage system. Animals used for Microglia isolation (5 ENR and 3 STD mice from each genotype) were killed through cervical dislocation.

Animals for histological analysis (28 ENR and 16 STD mice from each genotype) were killed with a mixture of ketamine/xylazine and a transcardial perfusion with 0.9% saline and 4% paraformaldehyde (PFA). Final blood samples were taken from the heart (right atrium) just before transcardial perfusion. Brains were left in 4% PFA over night at 4°C and were transferred to 30% sucrose before sectioning on a freezing microtome at 40 μm thickness.

### Analysis of RFID data

Radio-frequency identification (RFID) antennas, located in connecting tunnels, were used for longitudinal monitoring of behavioral activity within the cage system. Contact of mice to RFID antennas were recorded using the software PhenoSoft Control (PhenoSys GmbH, Berlin, Germany), which saved antenna and mouse identifiers together with the time stamp of the antenna contact into a database. Raw data of the antenna contacts reduced into 5s intervals and additional meta data are provided upon request. Data reduction and calculation of RE were performed as previously described^43, 44^. Because mice are nocturnal animals, only the events recorded during the dark phase were retained. Data from nights following events that could disturb patterns of exploration, such as cleaning of the cage or behavioral testing, were excluded. Cumulative RE was calculated by cumulative addition of mean RE from the 12 time-blocks.

### Social analysis

The social scores (social distance and following events) were calculated using the Rpackage ColonyRack (see https://rupertoverall.net/ColonyTrack/index.html for more details). In brief the social distance score is the average of the shortest distance within the cage system between the target mouse and all other mice. This is reported as an average for each mouse, for each day. A following event is recorded when one mouse triggers an antenna and another mouse triggers the same antenna within <1s, traveling in the same direction. The total number of events is reported for each mouse for each day.

### Analysis of blood samples and ELISA Protein detection

Blood was collected into EDTA-coated microvettes (Sarstedt). After 30 min, blood samples were centrifuges for 15 min at 2000 × g at room temperature. Plasma was collected from the supernatant, stored at -80°C. samples were sent to OLINK (Upsala, Sweden) and analyzed by proximity extension assay (PEA) for 96 proteins. Quality controls and data normalization was performed by Olink ^45^. Provided normalized protein expression (NPX) values on a log 2 scale were used for further group comparisons. The measured NPX values for all proteins are provided as Suppl. table 1). Blood samples taken at 7 months were procced for serum collection as described before were used to measure Casp3(E-El-M0238, Elabscience) and Ill1α (EA100140, Origene) protein levels by colorimetric sandwich ELISA following manufactures instruction.

### Histology

Tissue fixation and immunohistochemistry for the analysis of adult neurogenesis were performed as previously described (Ehret et al., 2020). Briefly, brains were cut into 40 µm coronal sections using a dry ice–cooled copper block on a sliding microtome (Leica, SM2000R). Sections were stored at 4°C in cryoprotectant solution (25% ethylene glycol and 25% glycerol in 0.1 M phosphate buffer). For the detection of plaques, every 12^th^ section of the brain was incubated in X-34 (dissolved in 60% PBS and 40% ethanol) for 20 min, followed by 3 quick washes in tap water and development in NaOH buffer (0.2 g% NaOH in 80% ethanol) for 2min. Subsequent sections were again washed in tap water for 10 min and transferred to PBS followed by counterstaining with 5 mM Draq5 (1:500, 65-0880-92, eBiosciences) for 1h before sections were mounted in 0.1 M phosphat buffer and cover-slipped with Flouromount G (Invitrogen) mounting media. Slides were stored at 4°C till analysis at the fluorescence microscope (Axio Imager.M2, Zeiss) using a HXP a 20 × Plan-Apochromat lens and the following filter settings for X-34 (FS49, with EX BP 365 and EM BP 445/50) for Draq5 (FS50, with EX BP 640/30 and EM 690/50) and equipped with motorized stage. A tile scan of the entire hippocampus was performed. The total surface covered by amyloid plaques was determined using a custom-written script based on the “Analyze particle” function of Fiji (National Institutes of Health; http://fiji.sc/), after defining the hippocampus as region of interest. The total surface occupied by plaques was then reported and related to the hippocampal area of each section. The following functions were applied: z-projection, maximum filtration with radius of 2pixel, automatic thresholding using *“*Max. Entropy*”* and particle analysis with a circularity between 0.5 and 1.00. Automatic detection was verified manually by a blinded second investigator.

Fluorescent staining of Iba1, Ki67, Trem2 and Draq5 was performed to detect and phenotype microglia in every 12^th^ section of the brain. After 10mM citric acid pretreatment (pH 6 + 0.05% Tween-20) for 20 min at 90°C and 10 min cool down, sections were washed in PBS 3 times. After blocking with 10% donkey serum and 0.2% Triton-X, sections were incubated with primary antibody of Iba1 (1:800, 019-19741,Wako), Trem2 (1:200, AF1729, R&D), Ki67 (1:400, 14-5698-82, eBioscience) over 48 h at 4°C. Afterwards, sections were extensively washed with PBS containing 0.1% Tween and incubated with secondary antibody cocktail containing Cy3 (713-165-147, Dianova) Dylight 405 (712-475-153, Jackson ImmunoResearch) and Alexa Flour 488 (711-545-152, Dianova) for 2h at room temperature. Followed by washing wit PBS, incubation with Draq5 (1:500, 62251, Invitrogen) for 10 min and cover slipped using Flouromount G and high precision coverslips. Sections were analyzed with Spinning Disk (Zeiss Axio Observer.Z1 using a 20x/0.8 Plan-Apochromat objective, Colibri LED for excitation at 365, 470, 590, 626nm) running tile scans by outlining the hippocampus and cortex area. A Z-stack of 4 images with 1 µm separation were used to a quire a stack of 3µm range. Image segmentation was performed using a Fiji customized macro to quantify the number of microglia, cell size, soma size, morphological changes using M-score (based on ^46^), proliferation rate (Ki67) and activation state (Trem2). The developed macro is a multi-step algorithm that segments nuclei (Draq5), microglia (Iba1), their soma, and identifies double-positive microglia expressing proliferation (Ki67) and activation (Trem2). As commonly done in image analysis, we initiated by pre-processing the images and applying a background subtraction using rolling ball algorithms with specific radii (50 for Draq5, 10 for Iba1, 40 for Ki67, and 10 for Trem2). We then applied four global thresholds (Default, Li, Triangle, and Moments) to segment the Draq5, Iba1, Ki67, and Trem2 markers, respectively. To distinguish somas from processes, we used the MorphoLibJ (v1.3.1) integrated library and plugin^47^ as previously done ^48^. This involved applying an opening morphological filter with a 4-pixel radius octagon. The area and circularity for microglia and their somas were quantified by using analyse particles function in Fiji. Information about area and circularity of microglia was utilized to calculate the M-score for each cell, following the formula described previously ^46^. Somas were defined as objects within the area range of 35 and 500 μm², while disregarding those not double-positive for Draq5 as nuclei marker. The overlay with segmented nuclei was achieved by using the Morphological Reconstruction from MorphoLibJ, with the segmented nuclei image as marker and segmented soma image as mask (connectivity = 8 as default). This process was repeated for the Ki67 image, first with the nuclei, as this proliferation marker is located within the cell nuclei, and then with the microglia soma image to determine the number of proliferating microglia. The threshold for Ki67 was set within a 30 to 100 μm² area range and circularity 0.28 - 1.00. This script was validated on a sample data set. Validation of automatic segmentation of Iba1 using Bland-Altman blot (Suppl. Fig 2). A small offset can be detected towards lower cell counts by automatic segmentation due to cells within dense activation clusters, but the difference was less than 30%.

For the detection of IdU^+^ cells, the peroxidase method was applied. Briefly, free-floating sections were incubated in 0.6% hydrogen peroxide for 30 min to inhibit endogenous peroxidase activity. For antigen retrieval, sections were incubated in prewarmed 2.5 M hydrochloric acid for 30 min at 37°C, followed by extensive washes. Unspecific binding sites were blocked in tris-buffered saline (TBS) supplemented with 10% donkey serum (Jackson ImmunoResearch Labs) and 0.2% Triton X-100 (Carl Roth) for 1 h at room temperature. Primary antibodies were applied overnight at 4°C (monoclonal anti–IdU 1:4000; SAB3701448, Merck). Sections were incubated in biotinylated secondary antibodies (Jackson ImmunoResearch Labs) for 2h at room temperature. Antibodies were diluted in TBS supplemented with 3% donkey serum and 0.2% Triton. Detection was performed using the VECTASTAIN Elite ABC Reagent (9 µg/ml of each component: Vector Laboratories, LINARIS) with diaminobenzidine (0.075 mg/ml; Sigma-Aldrich). All washing steps were performed in TBS. Sections were afterwards mounted onto glass slides, cleared with Neo-Clear (Millipore), and cover-slipped using Neo-Mount (Millipore). IdU+ cells were counted on every 6^th^ section along the entire rostro-caudal axis of the dentate gyrus using a bright-field microscope (Leica DM 750). The quantification was performed with a blinded investigator.

For doublecortin (DCX) detection the peroxidase method was applied as well with a similar procedure but without any antigen retrieval step. The following primary antibody (rabbit anti-DCX 1:500, ab18723, Abcam) and biotinylated secondary antibody (donkey anti-rabbit,1:750, Jackson Immuno) was used. Analysis was performed similarly as described before for IdU.

### sc-RNA-seq data analysis

Mapping and counting were done with Cell Ranger (https://10xgenomics.com). To build the reference, the mouse genome (mm10) as well as gene annotation (Ensembl 98) were downloaded from Ensembl. The reference was created with cellranger mkref (v5.01) as similarly to what is described by 10x (https://support.10xgenomics.com/single-cell-gene-expression/software/release-notes/build#mm10_2020A), with the difference that “pseudogene” was added to the allowed biotype patterns. The raw sequencing data was then processed with the ‘cellranger count’ command using Cell Ranger (6.0.1). Next, demultiplexing of individual samples was performed following the Seurat workflow “Demultiplexing with hashtag oligos (HTOs)” as implemented in https://github.com/ktrns/scrnaseq. The resulting count matrices were further processed using scanpy 1.8.1 ^49^. Only cells with at least 2000 counts, 900 detected genes, at most 60% of the counts in the top 50 genes, and at most 15% of the reads in mitochondrial genes. Normalization was done with ‘pp.normalize_total’. The top 5000 highly variable genes were detected with ‘ pp.highly_variable_genes’ (n_top_genes=5000). 50 principal components were computed on the 5000 highly variable genes. Data integration was performed with harmony using the two 10x libraries as batch key ^50^. A neighborhood graph was computed with ‘pp.neighbors’. A umap visulisation was computed with ‘tl.umap’. An initial clustering was computed with ‘tl.leiden’ (resolution=0.1). One of the clusters showed the distinct expression of rather macrophage specific genes (Apoe, Cybb, Mrc1, CD74, Lyz2, Pf4) and missing microglia specific genes (P2Ry12, Sparc, Tmem119), which we removed from further investigations. Next, computation of highly variable genes, principal component analysis, data integration, neighborhood graph and UMAP computation were repeated on the remaining cells. A new leiden clustering with resolution = 0.7 resulted in 8 clusters. Marker genes were computed with ‘tl.rank_genes_groups’. Trajectory inference was performed with ‘tl.pagà ^51^ and plotted with ‘pl.paga_compar’ (threshold=0.1). Differentially expressed genes between STD and ENR housing conditions were computed with the Wilcoxon rank-sum test and pooling the cells from all available samples. The false discovery rate was computed using the Benjamini-Hochberg procedure. The source code for the analysis is available from the authors upon reasonable request. The software stack for the analysis is available as a singularity container with tag b124ed0 under https://gitlab.hrz.tu-chemnitz.de/dcgc-bfx/singularity/singularity-single-cell/container_registry/11.

### Statistics

All experiments were carried out with the experimenter blinded regarding the experimental group. Statistical analyses were done using the statistical software Prism 9 (Graphpad) and R (R Core Team, 2014). Two-way ANOVA with Bonferroni posthoc analysis was applied to identify effects through housing and genotype using Prism and R.

To compare variance between groups, Brown-Forsythe test from the car package was used. Data were visualized using dot plot function with mean +/- standard error of mean(SEM), Box-Plots in Prism and the ggplot2 package in R ^52^.

To illustrate the social network the close contacts matrix was created using igraph Rpackage and social distances measures ^53^. To only isolate the close social connections, the mean group social distance was calculated and used as a cutoff point for igraph visualization.

To depict overlap in protein regulation, a Venn diagram was depicted using VennDiagram package in R.

For interaction analysis on multiple parameters, spearman’s rank order coeffects was calculated and plotted using cor function in R or Prism, with confidence intervals of 95%. To compare interaction between two parameters that were normally distributed, Pearsons R was calculated with Prism and for not formally distributes values Spearman’s R values are reported in the graphs.

FACS results are depicted as bar graphs with mean +/- SEM. Populations of single-cell sequencing are depicted as violin-blot (median is visualized as a line) or as stacked bar graph with mean + SEM.

## Acknowledgement

We are particular grateful to Takashi Saito, RIKEN Brain Science Institute, for providing us with the APP knock-in models used in this study. G. Kempermann recived funding from the Alzheimer Forschung Initative (#19038). We thank A. Karasinsky and S. Guenther for mouse handling and support in all animal experiments. We thank D. Severinov for support on data visualization. We are grateful for support from D. Lasse, C. Steinhauer and D. Glaeser on tissue processing and histology and all other members of the Kempermann laboratory for assistance and support and legal applications. Further, we thank S. White at the Imaging Platform of the DZNE Dresden for support and assistance.

## REFFERENCES

1. Livingston, G., et al. Dementia prevention, intervention, and care: 2020 report of the Lancet Commission. Lancet 396, 413–446 (2020).

2. Buchman, A.S., et al. Total daily physical activity and the risk of AD and cognitive decline in older adults. Neurology 78, 1323–1329 (2012).

3. Donovan, N.J., et al. Association of Higher Cortical Amyloid Burden With Loneliness in Cognitively Normal Older Adults. JAMA Psychiatry 73, 1230–1237 (2016).

4. Ehret, F., et al. Presymptomatic Reduction of Individuality in the App(NL-F) Knockin Model of Alzheimer’s Disease. Biol Psychiatry 94, 721–731 (2023).

5. Samtani, S., et al. Associations between social connections and cognition: a global collaborative individual participant data meta-analysis. Lancet Healthy Longev 3, e740–e753 (2022).

6. Fleck, J.I., et al. Distinct Functional Connectivity Patterns Are Associated With Social and Cognitive Lifestyle Factors: Pathways to Cognitive Reserve. Front Aging Neurosci 11, 310 (2019).

7. Lopes, J.B., Malz, M., Senko, A.N., Zocher, S. & Kempermann, G. Loss of individualized behavioral trajectories in adult neurogenesis-deficient cyclin D2 knockout mice. Hippocampus 33, 360–372 (2023).

8. Garthe, A., Behr, J. & Kempermann, G. Adult-generated hippocampal neurons allow the flexible use of spatially precise learning strategies. PLoS One 4, e5464 (2009).

9. Shors, T.J., et al. Neurogenesis in the adult is involved in the formation of trace memories. Nature 410, 372–376 (2001).

10. Snyder, J.S., Hong, N.S., McDonald, R.J. & Wojtowicz, J.M. A role for adult neurogenesis in spatial long-term memory. Neuroscience 130, 843–852 (2005).

11. Kempermann, G. Environmental enrichment, new neurons and the neurobiology of individuality. Nat Rev Neurosci 20, 235–245 (2019).

12. Kempermann, G., et al. The individuality paradigm: Automated longitudinal activity tracking of large cohorts of genetically identical mice in an enriched environment. Neurobiol Dis 175, 105916 (2022).

13. Bis, J.C., et al. Whole exome sequencing study identifies novel rare and common Alzheimer’s-Associated variants involved in immune response and transcriptional regulation. Mol Psychiatry 25, 1859–1875 (2020).

14. Pimenova, A.A., Raj, T. & Goate, A.M. Untangling Genetic Risk for Alzheimer’s Disease. Biol Psychiatry 83, 300–310 (2018).

15. Wightman, D.P., et al. A genome-wide association study with 1,126,563 individuals identifies new risk loci for Alzheimer’s disease. Nat Genet 53, 1276–1282 (2021).

16. Reemst, K., Noctor, S.C., Lucassen, P.J. & Hol, E.M. The Indispensable Roles of Microglia and Astrocytes during Brain Development. Front Hum Neurosci 10, 566 (2016).

17. Deerhake, M.E. & Shinohara, M.L. Emerging roles of Dectin-1 in noninfectious settings and in the CNS. Trends Immunol 42, 891–903 (2021).

18. Keren-Shaul, H., et al. A Unique Microglia Type Associated with Restricting Development of Alzheimer’s Disease. Cell 169, 1276–1290 e1217 (2017).

19. Wang, S., et al. Anti-human TREM2 induces microglia proliferation and reduces pathology in an Alzheimer’s disease model. J Exp Med 217 (2020).

20. Yi, Y.S., et al. p204 Is Required for Canonical Lipopolysaccharide-induced TLR4 Signaling in Mice. EBioMedicine 29, 78–91 (2018).

21. Zhao, H., et al. The roles of interferon-inducible p200 family members IFI16 and p204 in innate immune responses, cell differentiation and proliferation. Genes Dis 2, 46–56 (2015).

22. Kosel, F., Torres Munoz, P., Yang, J.R., Wong, A.A. & Franklin, T.B. Age-related changes in social behaviours in the 5xFAD mouse model of Alzheimer’s disease. Behav Brain Res 362, 160–172 (2019).

23. Muntsant, A., Castillo-Ruiz, M.D.M. & Gimenez-Llort, L. Survival Bias, Non-Lineal Behavioral and Cortico-Limbic Neuropathological Signatures in 3xTg-AD Mice for Alzheimer’s Disease from Premorbid to Advanced Stages and Compared to Normal Aging. Int J Mol Sci 24 (2023).

24. Locci, A., et al. Comparison of memory, affective behavior, and neuropathology in APP(NLGF) knock-in mice to 5xFAD and APP/PS1 mice. Behav Brain Res 404, 113192 (2021).

25. Misrani, A., et al. Mitochondrial Deficits With Neural and Social Damage in Early-Stage Alzheimer’s Disease Model Mice. Front Aging Neurosci 13, 748388 (2021).

26. Samaey, C., Schreurs, A., Stroobants, S. & Balschun, D. Early Cognitive and Behavioral Deficits in Mouse Models for Tauopathy and Alzheimer’s Disease. Front Aging Neurosci 11, 335 (2019).

27. Doludda, B., Bogado Lopes, J., Zocher, S., Kempermann, G. & Overall, R.W. Early-life experience determines the social stability of adult communities in female mice. bioRxiv (2023).

28. Kozorovitskiy, Y. & Gould, E. Dominance hierarchy influences adult neurogenesis in the dentate gyrus. J Neurosci 24, 6755–6759 (2004).

29. Maras, P.M., et al. Differences in microglia morphological profiles reflect divergent emotional temperaments: insights from a selective breeding model. Transl Psychiatry 12, 105 (2022).

30. Goldman, D.H., et al. Age-associated suppression of exploratory activity during sickness is linked to meningeal lymphatic dysfunction and microglia activation. Nat Aging 2, 704–713 (2022).

31. Koca, S., et al. Decreased levels of cytokines implicate altered immune response in plasma of moderate-stage Alzheimer’s disease patients. Neurosci Lett 786, 136799 (2022).

32. Zhang, H., Zhang, L., Zhou, D.M., Li, H.F. & Xu, Y. ErbB4 mediates amyloid β-induced neurotoxicity through JNK/tau pathway activation: Implications for Alzheimer’s disease. Journal of Comparative Neurology 529, 3497–3512 (2021).

33. Sierra, A., et al. Microglia shape adult hippocampal neurogenesis through apoptosis-coupled phagocytosis. Cell Stem Cell 7, 483–495 (2010).

34. Ekdahl, C.T., Claasen, J.H., Bonde, S., Kokaia, Z. & Lindvall, O. Inflammation is detrimental for neurogenesis in adult brain. Proc Natl Acad Sci U S A 100, 13632–13637 (2003).

35. Aloisi, F., Ria, F. & Adorini, L. Regulation of T-cell responses by CNS antigen-presenting cells: different roles for microglia and astrocytes. Immunol Today 21, 141–147 (2000).

36. Chen, W.T., et al. Spatial Transcriptomics and In Situ Sequencing to Study Alzheimer’s Disease. Cell 182, 976–991 e919 (2020).

37. Park, J., Baik, S.H., Mook-Jung, I., Irimia, D. & Cho, H. Mimicry of Central-Peripheral Immunity in Alzheimer’s Disease and Discovery of Neurodegenerative Roles in Neutrophil. Front Immunol 10, 2231 (2019).

38. Moore, Z., Mobilio, F., Walker, F.R., Taylor, J.M. & Crack, P.J. Abrogation of type-I interferon signalling alters the microglial response to Abeta(1-42). Sci Rep 10, 3153 (2020).

39. Heneka, M.T., et al. Neuroinflammation in Alzheimer’s disease. Lancet Neurol 14, 388–405 (2015).

40. Sanford, S.A.I. & McEwan, W.A. Type-I Interferons in Alzheimer’s Disease and Other Tauopathies. Front Cell Neurosci 16, 949340 (2022).

41. Bialas, A.R., et al. Microglia-dependent synapse loss in type I interferon-mediated lupus. Nature 546, 539–543 (2017).

42. Saito, T., et al. Single App knock-in mouse models of Alzheimer’s disease. Nat Neurosci 17, 661–663 (2014).

43. Zocher, S., et al. Early-life environmental enrichment generates persistent individualized behavior in mice. Sci Adv 6, eabb1478 (2020).

44. Freund, J., et al. Emergence of individuality in genetically identical mice. Science 340, 756–759 (2013).

45. Assarsson, E., et al. Homogenous 96-plex PEA immunoassay exhibiting high sensitivity, specificity, and excellent scalability. PLoS One 9, e95192 (2014).

46. Waller, R., et al. Iba-1-/CD68+ microglia are a prominent feature of age-associated deep subcortical white matter lesions. PLoS One 14, e0210888 (2019).

47. Legland, D., Arganda-Carreras, I. & Andrey, P. MorphoLibJ: integrated library and plugins for mathematical morphology with ImageJ. Bioinformatics 32, 3532–3534 (2016).

48. Davis, B.M., Salinas-Navarro, M., Cordeiro, M.F., Moons, L. & De Groef, L. Characterizing microglia activation: a spatial statistics approach to maximize information extraction. Sci Rep 7, 1576 (2017).

49. Wolf, F.A., Angerer, P. & Theis, F.J. SCANPY: large-scale single-cell gene expression data analysis. Genome Biol 19, 15 (2018).

50. Korsunsky, I., et al. Fast, sensitive and accurate integration of single-cell data with Harmony. Nat Methods 16, 1289–1296 (2019).

51. Wolf, F.A., et al. PAGA: graph abstraction reconciles clustering with trajectory inference through a topology preserving map of single cells. Genome Biology 20 (2019).

52. Wickham, H. ggplot2. Wires Comput Stat 3, 180–185 (2011).

53. Csárdi, G. & Nepusz, T. The igraph software package for complex network research. InterJournal **Complex Systems**, 1695 (2006).

