## Supplementary material for "Exploring social modulation: Microglia as a key mediator of individual immune response, plasticity and pathology in App-NL-G-F mouse model of Alzheimer’s disease": Suppl. 1-2

SUPPLEMENT:

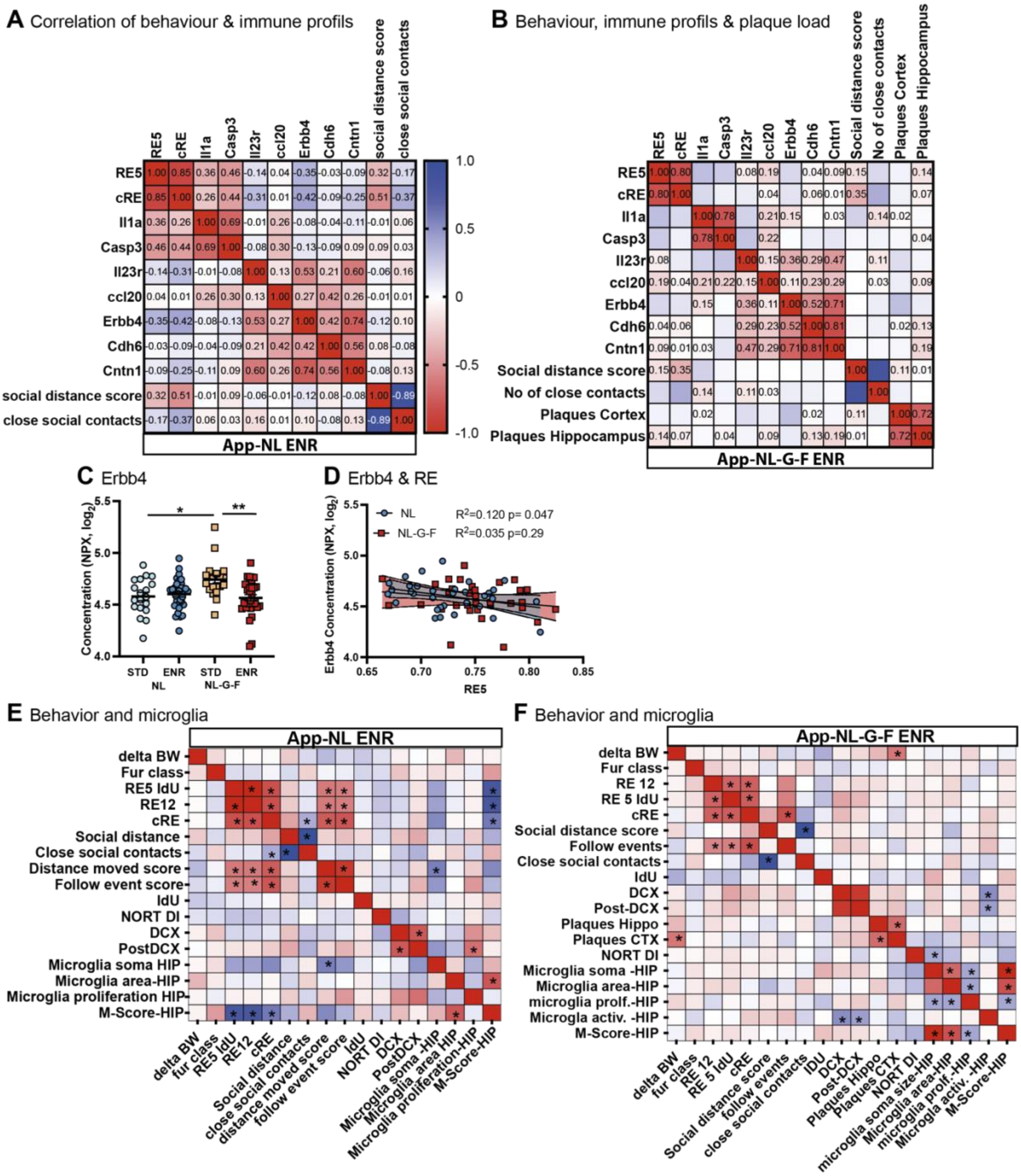


**Supplement 1: Analysis of correlative effects of social and explorative behavior, peripheral immune response and microglia state.**

(**A-B**) Correlation between behavior and social parameters and selected peripheral immune markers analyzed by proximity extension assay (PEA) at 3 months in NL (**A**) and NL-G-F (**B**) mice, Spearman's R values are color coded and written within relevant cells. (**C**) PEA analysis of Erbb4 in plasma at 3 months depicts effect of genotype and enrichment. (**D**) Correlation between Erbb4 concentration in plasma and explorative behavior measure by RE within the same time frame showed a significant positive effect in NL but not NL-G-F with a similar slope. Linear regression values are shown. (**E-F**) Behavioral correlation of NL (**E**) and NL-G-F (**F**) mice housed in enrichment. Analyzed by Spearmann's R, however correlation strength are not depicted due to space constrains but significant effects are indicated by*. Significant interactions are shown as * p<0.05 and**p<0.01.

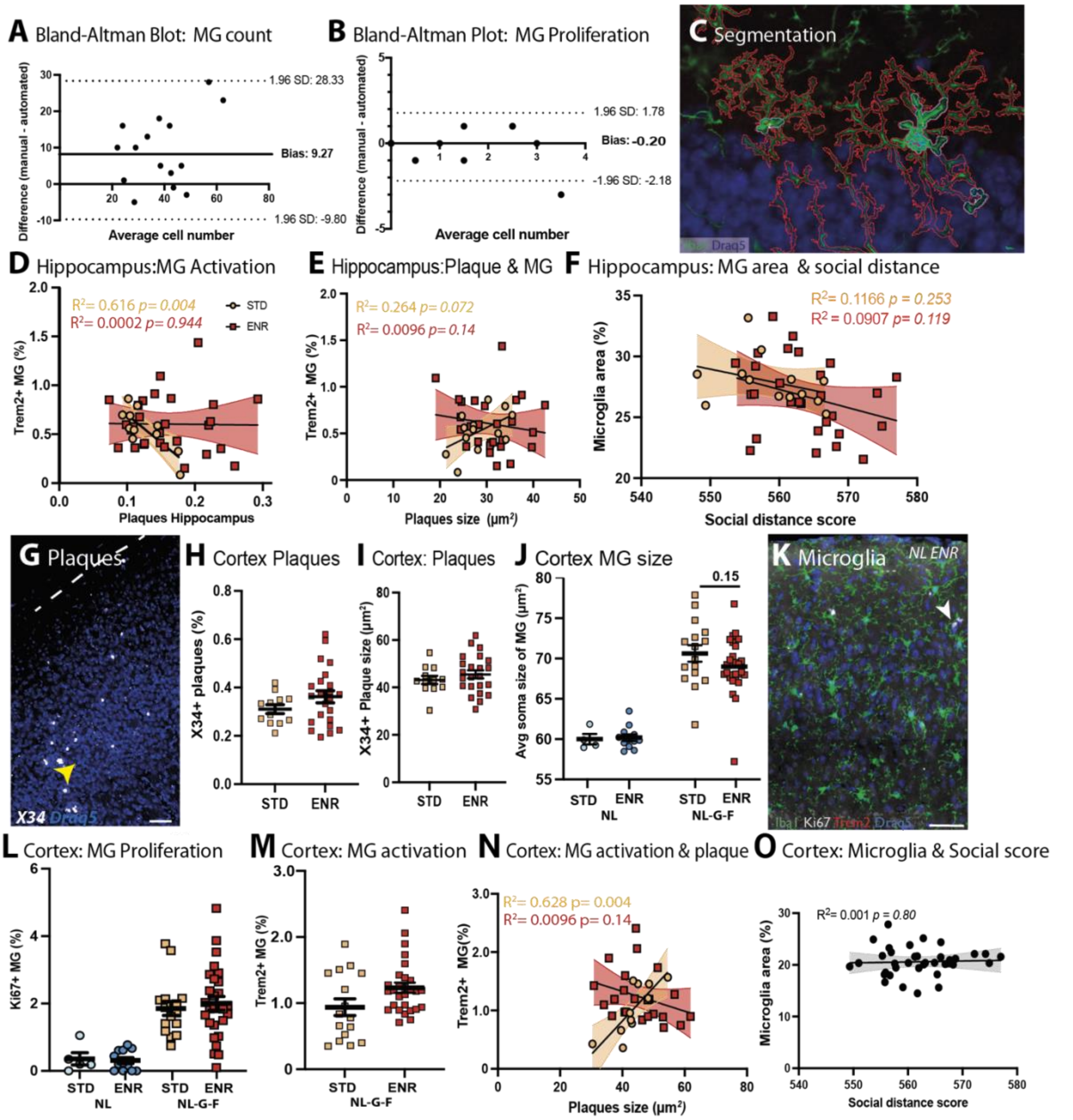


**Supplement 2: Semi-automated histological analysis of microglia in hippocampus and cortex**

(**A-B**) Bland-Altman blot comparing the sensitivity and accuracy of automated image segmentation vs. manual counting of Iba1+ microglia (MG) cells and Ki67+ MG. (**C**) Micrograph of MG (Iba1) with outlined segmentation of entire Microglia in red and soma in cyan. (**D**) The percentage of Trem2 dependent MG activation in hippocampus was analyzed against plaque load. **(E)**Trem2 dependent activation was analyzed against plaque size in the hippocampus**.** (**F**) Microglia area is in relation to social distance score here analysis of each genotype independently is shown to depict the similarity in slope. (**G**) Representative micrograph of plaques in the cortex labeled by X34. (**H-I**) Analysis of Plaque area (**H**) and plaque size (**I**) in the cortex did show any alterations due to ENR. (**J**)Average soma size of MG in cortex revealed that genotype has a major impact on soma size two-way ANOVA $F_{(1, 57)}$= 63.70, *p<0.0001;* with a trend of ENR to reduce soma size in NL-G-F p=0.13. (**K**) Representative micrograph of MG in cerebral cortex of control mice (NL) with no activation but proliferation of MG, using Iba1 (MG), Ki67 (Proliferation), Trem2 (Activation) and Draq5 (nuclei). Proliferating MG are indicated by white arrow. (**L**) Analysis of MG proliferation in cortex, showed increase proliferation in NL-G-F but no ENR effect but an increase in variance upon ENR was using Brown-Forsythe test *p = 0.035*. (**M**) MG activation was not altered upon ENR in cortex, however, a correlation to plaque size can be seen (**N**). (**O**) In the cortex no correlation of microglia area and social behavior can be found.
